# A regulated environment for micro-organs defines essential conditions for intercellular Ca^2+^ waves

**DOI:** 10.1101/081869

**Authors:** Cody E. Narciso, Nicholas M. Contento, Thomas J. Storey, David J. Hoelzle, Jeremiah J. Zartman

## Abstract

The mechanical stress state of an organ is a critical, but still poorly understood, driver of organogenesis and regeneration. Here we report a chip-based regulated environment for micro-organs (REM-Chip) that enables systematic investigations of the crosstalk between an organ’s mechanical stress environment and biochemical signaling under a multitude of genetic and chemical perturbations. This method has enabled us to identify essential conditions for generating organ-scale intercellular calcium (Ca^2+^) waves (ICWs) in *Drosophila* wing imaginal discs that are also observed *in vivo*. Spontaneous ICWs require the presence of components in fly extract-based growth serum (FEX). Using the REM-Chip, we demonstrate that the release and not the initial application of mechanical compression is sufficient but not necessary to initiate ICWs. Further, the extent of the Ca^2+^ response is heterogeneous between discs and correlates with the degree of spontaneous ICWs activity in the pre-stress state. This system and method enable detailed examinations of the interplay between mechanical stress state, biochemical regulatory networks, and physiology in complex, hierarchically organized organ cultures.

## SIGNIFICANCE STATEMENT

We present a first-of-class microfluidic chip that can perturb a developing organ both chemically and mechanically. Here we advance the field of organogenesis by presenting precise mechanical perturbation methods for whole-organ explants. This enables researchers to systematically interrogate critical relationships between mechanical stress state and biochemical signaling. Our methods advance available modes of studying *Drosophila* wing disc development, a powerful model for examining pathways critical for human development. We elucidate the causative links between mechanical perturbations and ICWs. Spontaneous ICWs require fly extract (FEX), a growth serum derived from flies, to be included in the culture media. Further, we find that the extent of ICW response to mechanical perturbation is determined by the spontaneous ICW activity prior to stimulation.

Organs develop in a diverse landscape of signals. However, the connections between exogenous forces, gene expression, and signal transduction in organ culture are still poorly understood. The calcium ion (Ca^2+^) has been demonstrated as a universal second messenger that regulates and coordinates a diverse range of intracellular processes such as proliferation and morphogenesis (1). Dysregulation of Ca^2+^ signaling via genetic and epigenetic modifications has been implicated in human diseases including cardiomyopathies (2), cancer metastasis (3), and neurodegenerative disorders (4). Furthermore, Ca^2+^ signaling has been shown to act as a central signal integrator in stem cell proliferation and homeostasis (5). The ubiquity of Ca^2+^ signaling makes it difficult to identify causative mechanisms of Ca^2+^ regulation and function in a given biological context (1). In particular, the roles of Ca^2+^ in mechanical signal integration has not been systematically investigated in epithelial tissues, despite observations that extreme mechanical perturbations such as wounding (6) and gross mechanical deformations (7) excite intercellular Ca^2+^ transients. Analogous to Ca^2+^ signaling, there is compelling evidence that implicates mechanical stress as a regulator of the cell cycle, differentiation, and cell survival (8–11). However, testing hypotheses on the relationships between the Ca^2+^ signaling, genetic background and mechanical signaling in organs is technically challenging because of the inability to apply precise mechanical perturbations to a specimen with current organ culture practices.

Microsystems have been increasingly used to mechanically perturb individual cells and cell populations. Microsystem designs range from structures with mechanical linkages for microscale manipulation (12) to contained vessels with integrated actuators for *in situ* perturbation (13). Microfluidic devices are an important class of microsystem that have been used to control chemical perfusion (14) and temperature (15) profiles. More recently, microfluidic devices have been used to mechanically perturb cells and confluent monolayers through deformation of microfluidic channel walls (16). In contrast to monolayers of a homogeneous cell type, organ-like cultures are particularly important for the study of development because they have innate morphogenetic diversity, hierarchical organization, and intact extracellular matrix (ECM). While organ-like cultures better replicate the *in vivo* environment, no device exists for combined mechanical manipulation and live-imaging of intact cultured organs (17). Here, we present a device designed for detailed mechanical manipulation of organ cultures.

The microsystem and method presented here is tuned to regulate the organ culture environment. Organ culture permits precise environmental perturbations with enhanced image quality compared to the *in vivo* context. However, organs are more difficult to culture and mechanically perturb than cells as they are generally less robust and less adherent to culture substrates. Our microfluidic device, termed the **R**egulated **E**nvironment for **M**icro-organs **Chip** (REM-Chip) is designed to test organs in an *ex vivo* context that closely mimics the *in vivo* microenvironment. The key features of the REM-Chip are: a gentle organ loading procedure; integrated fluidic channels to deliver growth media or other chemical constituents; deformable diaphragms to apply a compressive stress to an organ culture; and compatibility with small working distance objectives to enable real-time measurement of fluorescently labeled sensors.

To demonstrate the utility of the device, we used the *Drosophila* wing imaginal disc as a model organ. The wing disc has a powerful genetic toolkit and conserves many of the mechanisms of human development and disease (18). The wing disc develops in a naturally dynamic mechanical environment as larval motion is coupled to whole body deformation. Specifically, our group and others (7) have observed wing disc deformations during *in vivo* imaging of developing larva. In our studies, intercellular Ca^2+^ waves (ICWs) are observable in 34% of the discs (20 min of imaging, *n* = 37 and Wu et al., in preparation), but do not qualitatively correlate with physiological deformation during larval movement. In recent *in vivo* studies (7), mechanical compression could not be precisely prescribed, nor the Ca^2+^ response observed during the application of compressive stress. This device and method have enabled us to report that the release, and not the initiation of mechanical compression, stimulates an ICW that traverses the disc, elucidating the direct causative links between mechanical stress and ICWs. Spontaneous and mechanically-induced ICWs occur only if fly extract (FEX), a growth serum derived from flies, is included in the chemical milieu. Compression-release is sufficient to evoke ICWs under these conditions with the extent of ICW response determined by the pre-stress physiological state of the organ as measured by spontaneous ICW activity occurring in the stress free state. This highlights an important relationship between chemical and mechanical signaling revealed by our method. The REM-Chip and corresponding method is extensible to many other organ culture models and provides a powerful assay for measuring response to mechanical stimulation in organ growth, development, and homeostasis.

## RESULTS AND DISCUSSION

### REM-Chip system design

The REM-Chip is a two-layer microfluidic device developed for organ cultures that modulates the local chemical and mechanical microenvironment of a developing wing disc (Fig. 1). Both layers are made from the flexible, optically transparent, polymer polydimethylsiloxane (PDMS) and are bonded to a glass coverslip (Fig. 1A). Organ culture media and other chemical constituents flow in the bottom layer, and a pneumatically controlled deformable diaphragm in the top layer applies compression perpendicular to the imaging plane. The REM-Chip is integrated into a system composed of: a syringe pump for driving media flow; an array of pressure transducers to control diaphragm deflection; and a spinning-disc confocal microscope with coordinated stage translation and imaging routines (Fig. 1A, Supplementary Information, SI Fig. S1-3, **and SI text 1**). Earlier iterations of the REM-Chip design are shown in SI Fig. S4 and described in **SI text 2**.

**Fig. 1.**
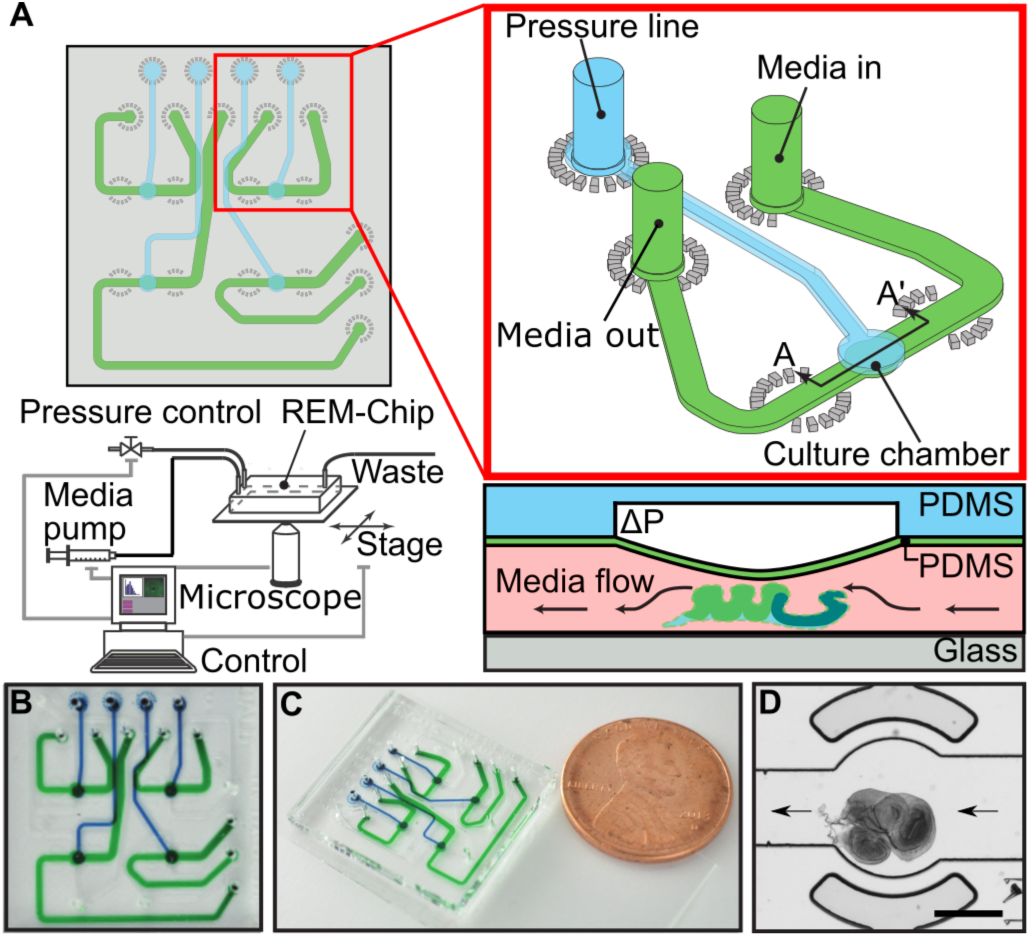
(**A**) REM-Chip schematic shown at top left with detail of a single chamber unit (red box). An individual culture chamber and accompanying fluid and pressure lines is integrated within a larger chip of four individually addressable culture chamber wells, for parallel experimentation, and part of a computer controlled network of external hardware (lower left). Diagram showing cross section of culture and pressure chambers through line A-A’ showing operation of the diaphragm mechanism for compressing wing discs. An applied pressure signal deforms a thin membrane in the chamber ceiling and compresses the disc below. (**B**, **C**) Photos of the device with individually addressable fluid (green) and pressure (blue) channels for each culture chamber. A US penny is shown for scale. (**D**) A 3^rd^ instar wing disc loaded into a culture chamber of a REM-Chip. Arrows indicate direction of media flow. Scale bar is 400 µm.

The wing disc loading protocol is optimized to reduce wing disc stress. The inlet channel of the REM-Chip is large enough (nominally 600 µm x 100 µm) for a wing disc to be loaded by flowing the disc through the channel. The channel is first flushed with organ culture media, and a wing disc is pipetted into the inlet. Next, the disc is positioned in the culture chamber by withdrawing organ culture media through the outlet channel with a pipette (**SI** Fig. S5 **and SI text 3**). Once loaded, culture media is flowed at 2 µL/h via a syringe pump to deliver media and chemicals; this low flowrate does not perturb the position of the wing disc. The area of the culture chamber is larger than an average wing disc to allow for variation in wing disc size and diametrical expansion during mechanical compression. The culture chamber size for wing discs is nominally 800 µm in diameter by 100 µm in height, but can be adjusted to accommodate other model systems.

### Validation of the REM-Chip

Compressive stress is applied to the wing disc by the REM-chip’s diaphragm, which is controlled via the pressure line. The deflection of the center of the circular diagram increases monotonically with applied pressure, as measured by confocal microscopy of diaphragms labeled by the fluorescent dye Nile red (Fig. 2A-B). A pressure in excess of 20 kPa results in contact between the diaphragm and the bottom of the chamber, setting the upper limit for compression experiments. The deflection-pressure relationship is in close agreement with a mechanics model of the deflection of a circular membrane under a uniform load (19) and is in agreement with the behavior of similar PDMS structures (20) (SI Fig. S6 and **SI text 4.2**). Deflection as a function of pressure for an individual diaphragm is repeatable to within ± 2 µm (s.d.), suggesting that variability in the system is negligible (Fig. 2B, SI Fig. S6A).

**Fig. 2.**
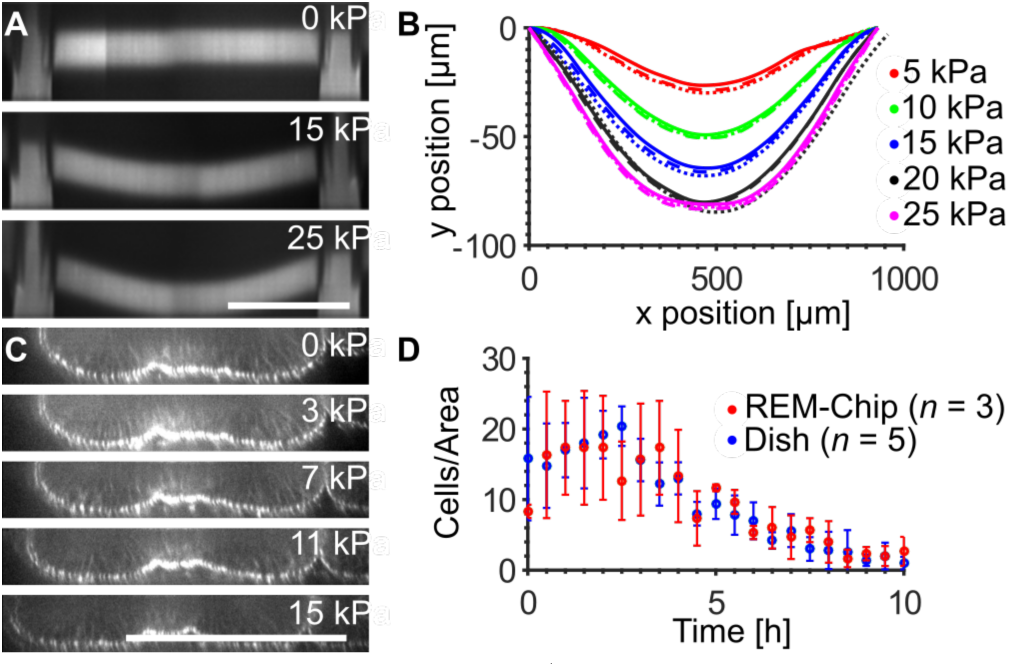
(**A**) Diaphragm stained with Nile red dye and imaged on a confocal microscope at indicated pressures. Scale bar, 400 um. (**B**) Measured diaphragm deflection profiles at the pressures indicated. Solid line denotes the 1^st^ applied pressure, dashed line denotes the 2^nd^ applied pressure, dotted line denotes the 3^rd^ applied pressure. Note the consistency of membrane position during each subsequent applied pressure. (**C**) Cross-section through the A-P axis of wing disc pouch showing deformation of the disc at the given applied pressures. Scale bar, 100 µm. (**D**) Proliferation curve. Cell divisions per 7656 µm^2^ area (250x250 pixels) per time point for wing discs cultured in the REM-Chip and in standard dish cultures. Sample size *n* = 5 for dish and *n* = 3 for REM-Chip, error bars are s.d. here and elsewhere.

Imaging of a wing disc compressed at increasing applied pressures demonstrates that the natural folds at the apical surface near the pouch flatten and expand when the disc is pressed against the glass coverslip by the diaphragm. This behavior is shown by a cross section through the center of a representative wing disc pouch shown in Fig. 2C.

Wing discs cultured in the REM-Chip using the optimized culture media formula WM1 (21) demonstrate no statistical difference in the number of mitotic cells when compared to discs cultured in a glass-bottom dish in the same media (repeated measures ANOVA, *p* = 0.95) (Fig. 2D). These results also agree with reported organ viability durations of ~12 hours on glass-bottomed wells (21, 22). Because culture in the REM-Chip does not significantly alter *ex vivo* disc viability relative to established methods, the REM-Chip can be used to study the effects of chemical and mechanical perturbations over the course of multiple hours.

### ICWs require a chemical signal

ICWs have been recently reported in wing discs both *in vivo* and *ex vivo* (7). We find that whole fly extract (FEX), a protein-rich serum often added to culture media to support growth and proliferation of *Drosophila* tissues, is necessary for the formation of ICWs *ex vivo* (Fig. 3 and Wu et al., in preparation). ICWs were first observed in *ex vivo* cultured wing discs cultured in WM1 media, which contains FEX (7). Removal of FEX serum from WM1 media results in a complete loss of ICW activity. Conversely, addition of FEX serum to ZB media, a chemically-defined culture medium, results in the formation of ICWs that are otherwise not present (Fig. 3). ZB media was engineered to support long-term proliferation and passaging of the wing-disc derived Cl.8 cell line (23). These results demonstrate that specific chemical stimulation by FEX is specifically required for the generation of both spontaneous and mechanically stimulated ICWs.

**Fig. 3.**
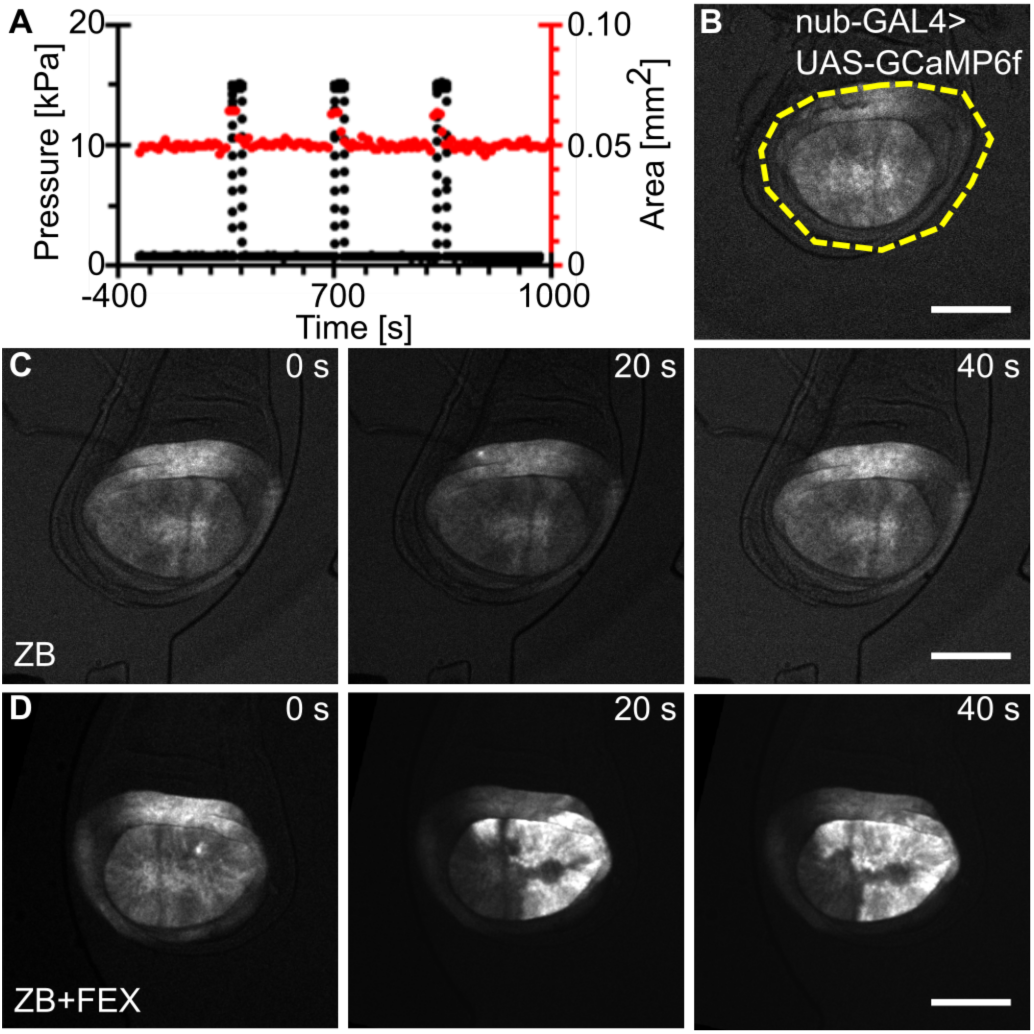
(**A**) Experimental plot of applied pressure with time and the change in pouch area with time for a cultured wing disc. Pulses in the pouch area (red line) correspond to pulses in the back pressure applied to the diaphragm (black line) and demonstrate mechanical perturbation of the wing disc. Time t=0 corresponds to the first pulse of compression. (**B**) Image showing the disc in the culture chamber prior to application of compression. *nub*-GAL4 drives expression of the genetically encoded Ca^2+^ sensor GCaMP6f to visualize intercellular Ca^2+^ signaling throughout imaging. The yellow dashed line indicates the maximum extent of pouch expansion during diaphragm deflection. (**C and D**) Montage of two discs cultured in ZB-based media with (**C**) or without (**D**) 5% FEX. Time 0 for each montage indicates the frame immediately following release of the first compression event. In the presence of FEX, compression release initiates an ICW (**C**). In the absence of FEX, intercellular transients are not observed following the release of compression (**D**). Scale bars are 100 µm.

### Release of compression initiates organ-wide ICWs

Mechanical perturbations have recently been implicated as a potential cause of the large ICWs in imaginal discs (7). However, experimental methods to allow wing disc observation during active compression or control the level and duration of the applied compressive stress were previously unavailable. Here, we have created a device that enables direct quantitative tests of the hypothesis that mechanical perturbation stimulates ICWs. A wing disc under a compressive stress applied by the diaphragm diametrically expands with a positive correlation with applied pressure (Fig. 3A-B). The release of this compressive stress initiates an ICW (Fig. 3C-D). ICWs are observed in 81% of compression events (*n* = 51/63 compressions) on release of compression in ZB media with 15% FEX. Compression arrests the formation of spontaneous ICWs. ICWs were never observed to formed during applied compressions for up to a 10-minute period (n=63 compressions). We observe 52% of discs (*n* = 11/21 discs) exhibit spontaneous ICW formation without mechanical compression within a 10-minute period. ICWs already underway were not stopped by application of compression (**SI Movie 9**).

Release of compressive stress exerted by a diaphragm backpressure of 15 kPa initiates an ICW approximately 20 s after release (Fig. 3C). Compression-release in the absence of FEX is not sufficient to initiate an ICW, indicating that mechanical initiation of ICW activity is dependent on the presence of FEX in the media (Fig. 3D). The outer folded region of the *nub*>*GCaMP6f* expressing domain of the wing disc is oriented such that the apical-basal direction of cells is in the plane of axial compression, resulting in lateral stretching in this region of the disc when the diaphragm is deflected. This portion of the wing disc shows persistent, active ICWs, suggesting that stretching promotes ongoing Ca^2+^ activity.

### Mechanically induced ICWs depend principally on the organ’s physiological status

To investigate important factors governing the extent of ICW activity, we performed a series of compression experiments varying duration and magnitude of compression over multiple biological experiments (*n* =21, 3 experiments). Each wing disc was loaded into the REM-Chip, held at 0 kPa back pressure for the first 10 min, and then subjected to a 10 kPa, 15 kPa, or 20 kPa diaphragm backpressures. A series of three compression/no-compression cycles were performed on each disc. The order of duration of compression was randomized: 30 s, 300 s, and 600 s. The wing disc response to mechanical compression-release events for this data set could be binned into two differing populations. Discs that showed qualitatively consistent spontaneous ICW activity in the first 10 minutes before compression consistently responded to mechanical compression-release stimulation (*n* = 34/36 compressions) at a global tissue level for all applied backpressure levels and durations (Fig. 4A-C). Discs that are naturally quiescent, exhibiting low-medium spontaneous ICW activity (small bursts) in the first 10 minutes of culture, responded to compression-release events much less frequently (*n* = 17/27) with low- to medium-ICW activity regardless of backpressure level or durations (Fig. 4A-C). In many cases, Ca^2+^ burst area due to compression-release covered the same spatial extent as the previous spontaneous pre-compression ICW events and more localized ICWs frequently initiated near the edge of the pouch with an inward direction towards the geometric center of the tissue. Significant population-level effects were observed in the peak mean pouch intensity after release (*p*<0.01), ICW burst time (*p*<0.001), and fraction of wing disc pouch occupied by the ICW (*p*<0.001). Student’s t-test with Bonferroni correction was used to obtain *p*-values.

**Fig. 4.**
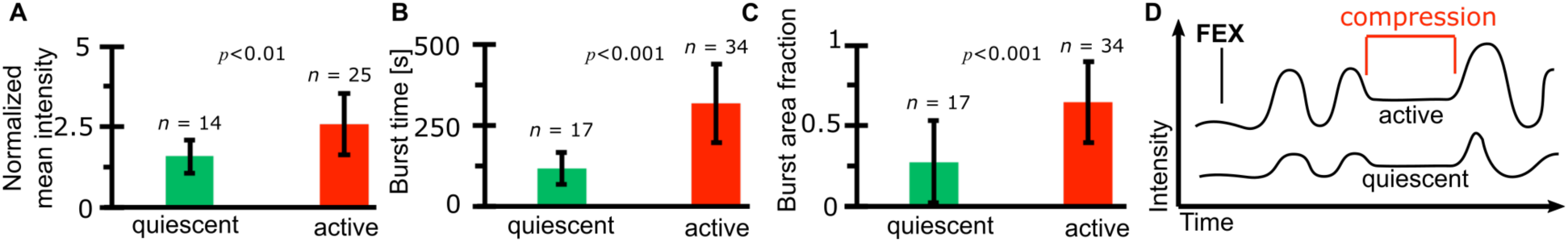
(**A**) Plot of the peak mean pouch intensity following release of compression normalized to the basal Ca^2+^ level. Active discs produce much larger ICWs (**C**) that tend to last longer (**B**), resulting in a significantly increased peak mean pouch intensity (*p* < 0.001). (**B**) Plot of ICW burst time calculated as the last frame that an ICW is observed following compression minus the first. Active discs produce ICWs that last longer, regardless of applied backpressure or hold times applied. (**C**) Plot of fraction of total pouch area occupied by the mechanically induced ICW. Active discs produce much larger ICWs than quiescent discs regardless of the applied pressure or hold time. (**D**) These results and observations lead to the working model that release of mechanical compression amplifies and forces the existing FEX induced ICW response. *p*-values are from a Student’s t-test with Bonferroni correction.

In comparison to the sub-population effect, we saw comparatively minor effects of applied backpressure, pouch deflection or backpressure hold time on the burst area, burst time, normalized mean peak intensity in the pouch, velocity or delay of the ICW response over the ranges tested. A small qualitative increase was observed in the effects of applied backpressure on burst time of the ICW. Additionally, there is a small qualitative increase in the max mean pouch intensity as a function of hold time and applied backpressure. These relationships are shown in SI Fig. S9. Compression-release induced ICW velocity was found to be 0.8 µm/s ± 0.6 µm/s, and the time to initiation was found to be 20 s ± 30 s (s.d.) after the compression-release. Taken together, the magnitude of compression-release induction of ICWs is dependent on the physiological state of the wing disc more than on duration or degree of deformation for the conditions tested (Fig. 4D).

### Mechanically stimulated ICWs result from IP_3_-mediated release from intracellular stores and depend on gap junction activity

We performed a targeted genetic screening approach to define the mechanism of ICW formation occurring due to mechanical stimulation compared to wound induced and spontaneous ICWs. IP_3_ signaling is highly conserved and present in both *Drosophila* and vertebrate models as an important mediator of Ca^2+^ release. The IP_3_ signaling cascade releases Ca^2+^ from internal stores in the endoplasmic reticulum (ER) by binding with its receptor, IP_3_R. Previously, we have delineated that laser-ablation induced intercellular Ca^2+^ flashes that Ca^2+^ transients in wing discs through IP_3_-mediated Ca^2+^ release and subsequent propagation through gap junctions (GJs) (24). This mechanism is well-established for intercellular transients as a result of wounding (6, 25) and is also responsible for waves observed during *Drosophila* oogenesis (26). To test whether mechanical induction of ICWs also rely on this conserved IP_3_-mediated mechanism, RNAi was used to knock down key components of this cascade (Fig. 5A). We found that inhibition of GJs through Inx2^RNAi^, inhibition of the IP_3_ receptor (IP_3_R^RNAi^), and inhibition of the endoplasmic reticulum (ER) pump SERCA (SERCA^RNAi^) all completely abolish the compression-release induced ICWs (Fig. 5B-C). Taken together, these results suggest that IP_3_-mediated release from intracellular stores and GJ communication between neighboring cells are necessary for the propagation of mechanically induced ICWs. Knockdown of SERCA shows a characteristic increase in basal Ca^2+^ level consistent with impairment of the ability to uptake cytoplasmic Ca^2+^ into the ER (Fig. 5C). This suggests that once the concentration of Ca^2+^ in the ER stores drop below a critical threshold, cells are unable to respond through this mechanism. This is in agreement with recently published work on wound-induced ICWs in *Drosophila* wing discs (7, 24). Interestingly, although GJ inhibited discs (Inx2^RNAi^) do not develop ICWs, increased flashing of individual cells is observed on release of compression (Fig. 5D-E). Cumulatively, these results suggest that compression-release increased Ca^2+^ spiking in individual cells, which was likely mediated by IP_3_. GJ activity enables transmission of IP_3_ and Ca^2+^ throughout the tissue. Additional details on all RNAi lines and their validation can be found in (**SI** Fig. S7 **and SI text 5**)

**Fig. 5.**
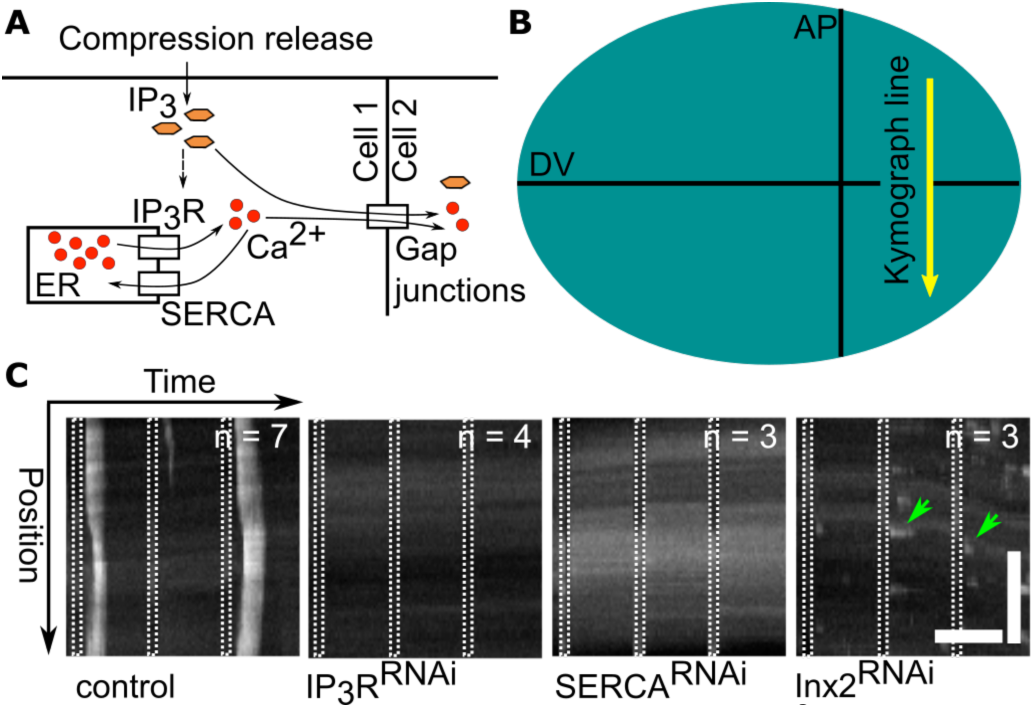
(**A**) Illustration of key Ca^2+^ signaling components involved in mechanically induced ICWs in the wing imaginal disc. (**B**) Schematic of the sampling line for the kymographs shown the wing disc in (**C**). (**C**) Intensity of GCaMP6f signal as a function of time along sampling line for representative samples from the indicated populations. Sample size for each population is indicated. White bands represent periods of compression. The control represents the tester line (*nub*-GAL4>UAS-*GCaMP6f*) and the gene of interest lines are each crossed to the tester line and are detailed in the (**SI text 5**). Taken together, this data strongly suggests that the ICW response is the result of IP_3_-mediated release from intracellular stores in the ER. Temporal scale bar (horizontal) represents 300 s, spatial scale bar (vertical) represents 100 µm. Although Inx2^RNAi^ does not exhibit an ICW as a result of compression, it does show an increase in stochastic single cell flashing (green arrows) following compression events. This indicates that although the disc is unable to respond by generating an ICW, it can still sense the applied perturbation.

### The REM-Chip enables new chemical and mechanical assays for organ cultures

Mounting evidence suggests that both chemical and mechanical cues impact organ development (27, 28). The REM-Chip is a powerful new tool for providing controlled mechanical and chemical microenvironments compatible with long-term organ cultures. Recent observations have linked mechanical stress to ICWs in wing discs (7); however, these experiments relied on manual methods to apply mechanical perturbations. These methods allow little control over magnitude or extent of the applied perturbation. Our REM-Chip design allows reproducible and controlled compressions of wing discs. Importantly, the ICW response to mechanical perturbation is found to be dependent specifically on the release, rather than the application, of compressive stress – a distinction lacking in previous observations of mechanically induced ICWs in both cultured cells and tissues. This observation is significant for two reasons. First, developing wing discs and other organs are constantly deformed via organismal and morphogenetic movements. Second, this observation posits interesting questions for further investigation. For instance, what are the downstream effects of mechanically induced ICWs on transcription, translation and mechanical properties of tissues? While other approaches have enabled mechanical perturbation of *Drosophila* systems (7, 12, 29, 30), the high fidelity mechanical control enabled by the REM-Chip expands the scope of available mechanical assays for organ culture. More generally, the REM-Chip system provides an attractive characterization tool to study the mechanotransduction properties of heterologous channels and mechanoreceptors that can easily be expressed using current *Drosophila* genetic tools. In addition the REM-Chip will enable future efforts that seek to develop and test novel pressure-based synthetic biology input/output modules that integrate both chemical and mechanical inputs.

### Chemical and genetic perturbations modulate mechanically stimulated ICWs

The REM-Chip has revealed that the release of compression, is a potent inducer of ICWs. ICWs were observed after mechanical compression only in the presence of FEX, indicating that both chemical and mechanical factors play a role in this response. This leads to two possibilities requiring future investigations: 1) either FEX is a permissive growth medium that enables intrinsic Ca^2+^ oscillations to occur in the wing disc coordinated through GJ communication; or 2) an unknown instructive component of FEX generates ICWs. Ongoing work (Wu et al. in preparation) favors the latter explanation.

RNAi against key components of the IP_3_ signaling cascade revealed that ICWs result from IP_3_-mediated release and travel through GJ-mediated communication. While Inx2^RNAi^ does not allow for the formation of ICWs, increased flashing of individual cells is observed on release of compression, suggesting that cells are still able to sense mechanical perturbations. This provokes the question of whether compressive stress is mainly inhibiting production of IP_3_ rather than closing GJs. While mechanical stimulation is sufficient to initiate ICWs, it is not the primary driver of ICWs, as they can be observed spontaneously in the absence of any observable mechanical perturbation. Rather a chemical signal contained in the FEX serum is the primary source of spontaneous ICWs in wing discs. The exact nature of such chemical factors and whether they may be developmentally regulated is a subject for future research.

### The REM-Chip enables long duration studies for a multitude of model organs

The REM-Chip does not inhibit the long-term viability of the wing disc in comparison to established *ex vivo* methods. Furthermore, chemical and mechanical perturbations are completely automated, paving a clear path for long-term investigation of mechanical and chemical perturbations on organ growth and homeostasis, as previously studied in cell cultures (31, 32), but not in organ cultures. Other potential long-term studies include determining how mechanical and chemical perturbations affect morphogenetic patterning in tissues. Lastly, although this specific REM-Chip detailed here is tailored for the *Drosophila* wing disc, simple modifications to channel and chamber dimensions will enable identical assays to be applied to other *Drosophila* organs in addition to model organs excised from *Xenopus*, zebrafish, and human organoids, to name a few.

## MATERIALS AND METHODS

### REM-Chip design and fabrication

REM-Chip layer designs were drafted in AutoCAD and each design was converted to a photolithography mask by printing on a transparency film (Fineline Imaging, Inc.). Microfluidic masters, composed of a silicon wafer and SU-8 photopolymer (SU-8 3050, MicroChem Corp.), were fabricated via standard lithographic micromachining methods (33). The microfluidic channels in the REM-Chip were defined using reverse micromolding of PDMS on the microfluidic masters–also termed soft lithography–and bonded together via standard methods for two-layer microfluidic devices (34, 35). Ports were punched in the PDMS device and the entire device was cleaned using isopropyl alcohol and then bonded to a glass coverslip (34). Complete details can be found in our published protocols(34). Completed REM-Chips were stored in covered petri dishes.

### Systems control and disc image acquisition

Media flow from the syringe reservoir to the REM-Chip devices was driven using a Harvard Apparatus Pump 11 Elite programmable syringe pump. Pressure signals to the REM-Chip were externally controlled by a custom-built pressure regulator box consisting of a manual regulator to step down the house air pressure, a bank of four electropneumatic pressure regulators (ITV001-3UML, SMC Corp., Tokyo, Japan) connected to the source pressure manifold, and an analog output module and accompanying hardware (NI 9264, National Instruments, Austin, TX) to electronically control the reference pressure. Each analog output channel was controlled using a custom-written LabVIEW (National Instruments, 2014) script (**SI text 1**).

Image acquisition was performed on a Nikon Eclipse T*i* confocal microscope (Nikon Instruments Inc., Melville, NY) with a Yokogawa spinning disc using an iXonEM+ cooled CCD camera (Andor Technology, South Windsor, CT). Image acquisition was controlled using MetaMorph v7.7.9 software (Molecular Devices, Sunnyvale, CA).

### Sample preparation

Wing discs were cultured in the chemically-defined, ZB-based media (23) or WM1 (21) media optimized for organ culture. Each culture chamber was prepared by filling a Becton Dickinson (BD) Plastipak 1 mL syringe with 1 mL of ZB-based or WM1 media, evacuating all air bubbles from the syringe, and pre-filling the microfluidic network with media. Culture chambers were visually inspected under a stereomicroscope to ensure no air bubbles were present in the culture chamber or fluid lines prior to wing disc loading. Imaginal discs from wandering third instar larvae were then dissected in the relevant culture media. Immediately following dissection, discs were loaded into prepared REM-Chip culture chambers by pipetting the disc over the fluid inlet in a small droplet of media. The disc was positioned with a dissecting needle such that the dorsal side was facing the inlet and the disc drawn into the culture chamber by manually withdrawing media from the fluid outlet with a pipette (SI Fig. S5). After all discs were loaded into culture chambers, the REM-Chip was affixed to the microscope stage, and fluid inlet lines were connected to the media reservoir and pre-flooded with culture media. Inlet lines were then plugged into the REM-Chip and media were perfused at a flow rate of 2 µL/h for the duration of all experiments.

### Diaphragm and wing disc deformation study

Diaphragm deflection was measured from images of a fluorescently labeled diaphragm over a range of applied pressures. The REM-Chip was first perfused with 1.5 mM Nile red dye (Santa Cruz Biotech) in methanol for at least 1 h to label all PDMS channel walls. Next, the Nile red was then evacuated from the culture chamber with deionized water; Nile red is insoluble in water. The diaphragm was imaged using confocal microscopy at applied pressures of 0, 5, 10, 15, 20, and 25 kPa. Confocal images were post-processed in Fiji (36) to stitch adjacent fields of view and obtain *xz* plane diaphragm images from the confocal z-stacks. Diaphragm deformation profiles were then measured using a custom MATLAB script that filters and thresholds the *xz* images, detects the diaphragm edges given a user-selected region of interest, and converts pixels to distance via microscope calibrations (see **SI text 4.1**).

Wing disc deformation under mechanical compression was studied via spinning disc confocal microscopy. *Drosophila* lines expressing DE-Cadherin::GFP (37), which concentrates in the subapical region of the wing disc and labels apical cell boundaries, were used to image the unbuckling and diametrical expansion of the pouch. Wing discs were excised, prepared, and loaded as per the standard protocol described above. Back pressure (0, 3, 7, 11, 15 kPa) was statically applied to the REM-Chip diaphragm and a confocal *z*-stack centered on the pouch was acquired at 60X magnification in 1 µm intervals. Each *z*-stack of images was then processed using Fiji/ImageJ (36) to generate a cross-sectional image along the AP axis of the wing disc.

### REM-Chip viability study

Flies expressing Lac::YFP, a green fluorescent marker for cell boundaries, were staged for 2 hours in vials at 25 **°**C. Larvae were collected for dissection at 120 hours (5 days) after egg laying. Discs were dissected and loaded into the REM-Chip in WM1 media as described above or dissected and loaded into glass bottom dishes for standard culture comparison as previously described (21). Discs were imaged at 40X magnification every 30 minutes for a minimum of 12 hours. A 250x250 pixel region, representing ¼ of the total frame size, was cropped from each video frame, centered on the wing disc pouch. The cropped regions were saved with descriptive filenames. The filenames were then randomized and a key automatically created using a batch script written by Jason Faulkner (**SIFiles.zip**) to create a blinded set of images. The numbers of mitotic cells per area were tabulated by counting the number of cells with increased apical area. The data were compared for statistical difference using a repeated measures ANOVA test.

### ICW mechanism study

A tester line was created by recombining *nub*-GAL4 with UAS-*GCaMP6f* on the second chromosome. This line was crossed with UAS-Gene-of-interest^RNAi^ lines to perturb essential components of the IP_3_ signaling cascade (full details provided in **SI text 5**). Discs from wandering third instar larvae from each cross were dissected and loaded into the REM-Chip as previously described. These discs were imaged at 10X magnification on a Nikon Eclipse T*i* confocal microscope for a total of 30 minutes, imaging every 10 seconds. While being imaged, the discs were subjected to the following schedule of applied pressures: 300 s at 0 kPa, 30 s at 15 kPa, 300 s at 0 kPa, 30 s at 15 kPa, 300 s at 0 kPa, 30 s at 15 kPa, 300 s at 0 kPa. Pressure fluctuations had a rise time of 1 sec. Videos were qualitatively scored for the presence or absence of a ICW on release of compression for each RNAi cross tested. The control group consisted of uncrossed discs from the tester line, subjected to the same schedule of applied pressures and imaging regiment.

### REM-Chip compression study

Wandering third instar wing discs from the tester line were dissected and loaded into the REM-Chip as previously described. Each experiment consisted of a set of three, unique, 3^rd^ instar wandering wing discs tested in parallel. Each wing disc was subjected to either 10 kPa, 15 kPa or 20 kPa diaphragm back pressure for a randomized set of three compression durations consisting of 30 s, 300 s, and 600 s each. Each compression period was preceded and followed by a 600 s quiescent period (0 kPa) to observe the Ca^2+^ dynamics. Discs were imaged as described above, except the total imaging period was extended to 60 minutes to accommodate the increased backpressure duration. Videos were randomly renamed as described above for blind analysis and the time to initiation, velocity, burst time, and burst area fraction and peak mean pouch intensity of each mechanically induced ICW were quantified using a combination of Fiji and MATLAB, complete details of the analysis are provided in (**SI text 6** and SI Fig. S8).

### Fly lines and reagents

Ca^2+^ signaling was visualized using the GAL4-UAS system (38). The Ca^2+^ signaling sensor UAS*-GCaMP6f* (39) (Bloomington stock #42747) was recombined with the *nubbin-*GAL4 driver (Bloomington stock #25754) to provide a readout of Ca^2+^ activity in the wing disc pouch (*nub*-GAL4>UAS-*GCaMP6f*), which we term the GCaMP6 tester line. For genetic perturbation experiments, the GCaMP6f tester line was crossed to TRiP lines for the targeted gene (all genotypes listed in **SI text 5**). All flies were raised at 25 °C, 70% relative humidity on cornmeal agar (40). All reagents are listed in (**SI text 5**).

### Image analysis routines

Supplemental movies to accompany figures are described in (**SI text 7**). All scripts used in the analysis of image data are included in (**SIFiles.zip**) and described in (**SI text 8**).

## ACKNOWLEDGEMENTS

The work in this manuscript was supported in part by NSF Award CBET-1403887 and the Notre Dame Advanced Diagnostics & Therapeutics Berry Fellowship. The authors gratefully acknowledge the Notre Dame Integrated Imaging Facility and the Notre Dame Nanofabrication Facility for the use of their imaging and fabrication facilities, respectively, Melinda Lake and Maxwell Kennard for their assistance with REM-Chip fabrication and pressure and vacuum regulator system design, respectively. The authors thank the Developmental Studies Hybridoma bank for antibodies. The authors thank S. Shvartsman, J. Boerckel, S. Restrepo and members of the Zartman lab for feedback on the manuscript.

## AUTHOR CONTRIBUTIONS

D.J.H. and J.J.Z. conceived the project. C.E.N. performed the organ culture studies. N.M.C. performed diaphragm deflection studies. T.J.S. participated in REM-Chip design including the pressure and vacuum regulator system and contributed to Fig. 2. D.J.H. initially designed REM-Chip and integrated system. C.E.N., N.M.C., D.J.H. and J.J.Z designed the experiments, analyzed the results and wrote the paper.

## Supplemental Information

### 1. Description of control system

Electropneumatic regulators (ITV0010-3UML and ITV-0090-3UML, SMC Pneumatics, Japan) were controlled with analog input (NI 9205, National Instruments, USA) and output (NI 9264) modules connected to a data acquisition card (NI eDAQ 9174). Pressure and vacuum lines were connected to the regulators following the manufacturer’s instructions. The input pressure was stepped down to <200 kPa using a manual pressure valve (4ZM08, Speedaire, USA). The system was powered by an NI PS-15 power module connected directly to wall power. Details of the different input and output channels are provide in Table S1. A diagram of the wired assembly for pressure and vacuum—vacuum was not used in this study—control is shown in Fig. S1. Photographs of the control system used in this study are shown in Fig. S2. The module IDs given in the table are important for properly controlling the hardware with the LabVIEW virtual instrument (VI) program (LabVIEW 2014 SP1) shown in Fig. S3. The full LabVIEW virtual instrument file (pressure_automation.vi) is available for download online.

#### Flow control materials

The fluid inlet lines (1/32” OD PEEK tubing, IDEX Health and Sciences, #1581) were attached to the syringe reservoir using a female-to-female luer (P-638), a headless nut (P-232), and a flat-bottomed ferrule (P-248).

**Table S1.** Description of input and output channels for the electropneumatic control system.

| Analog Input (NI 9205) |  |  |  | Analog Output (NI 9264) |  |  |  |
| --- | --- | --- | --- | --- | --- | --- | --- |
| Channel | Description | Module ID | Signal (V) | Channel | Description | Module ID | Signal (V) |
| A | Vacuum Output | AI19 | 1–5 | A | Vacuum Input | AO11 | 1–10 |
| B | Vacuum Output | AI18 | 1–5 | B | Vacuum Input | AO10 | 1–10 |
| C | Vacuum Output | AI17 | 1–5 | C | Vacuum Input | AO9 | 1–10 |
| D | Vacuum Output | AI16 | 1–5 | D | Vacuum Input | AO8 | 1–10 |
| E | Pressure Output | AI13 | 1–5 | E | Pressure Input | AO3 | 1–10 |
| F | Pressure Output | AI12 | 1–5 | F | Pressure Input | AO2 | 1–10 |
| G | Pressure Output | AI11 | 1–5 | G | Pressure Input | AO1 | 1–10 |
| H | Pressure Output | AI10 | 1–5 | H | Pressure Input | AO0 | 1–10 |
| GND | GND Terminal | COM, AISENSE |  |  |  |  |  |

**Figure S1.**
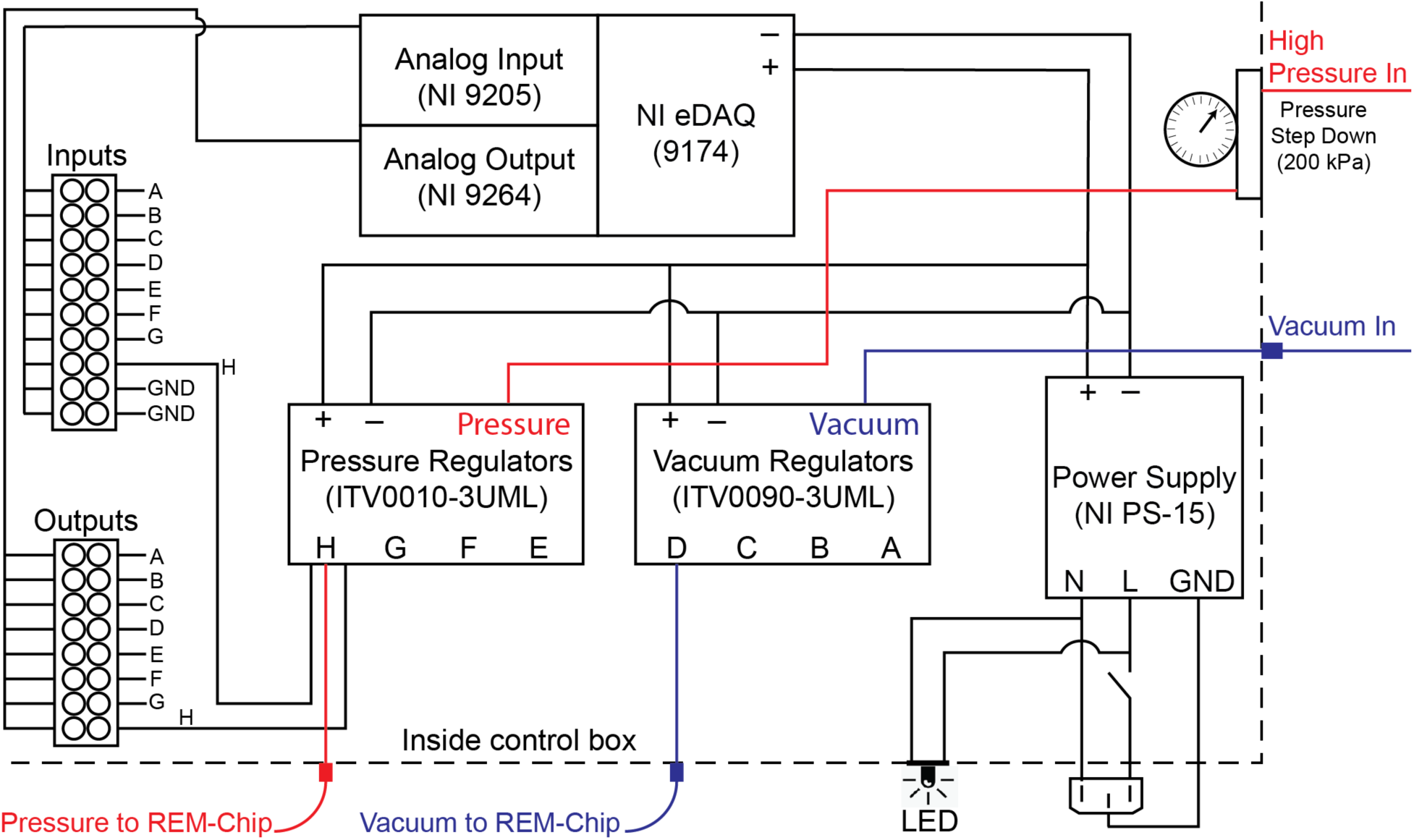
Control system wiring. Diagram of control system electrical wiring (black), pressure connections (red), and vacuum connections (blue). To avoid confusion, the wiring diagram is shown for one pressure regulator channel, and only one pressure and vacuum output are shown.

**Figure S2.**
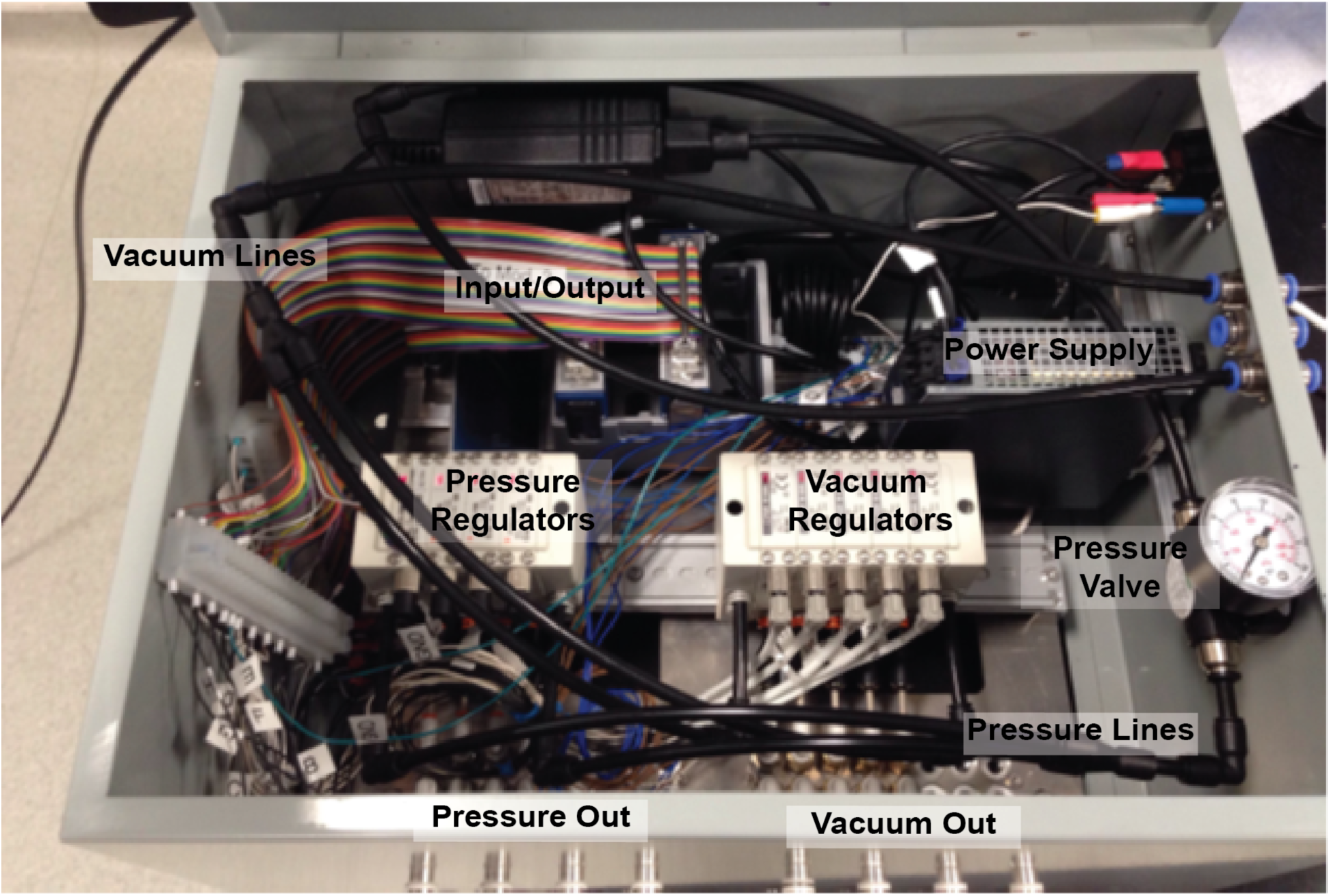
Control system. Photograph of the assembled control system used in this study corresponding to Fig. S1 schematic.

**Figure S3.**
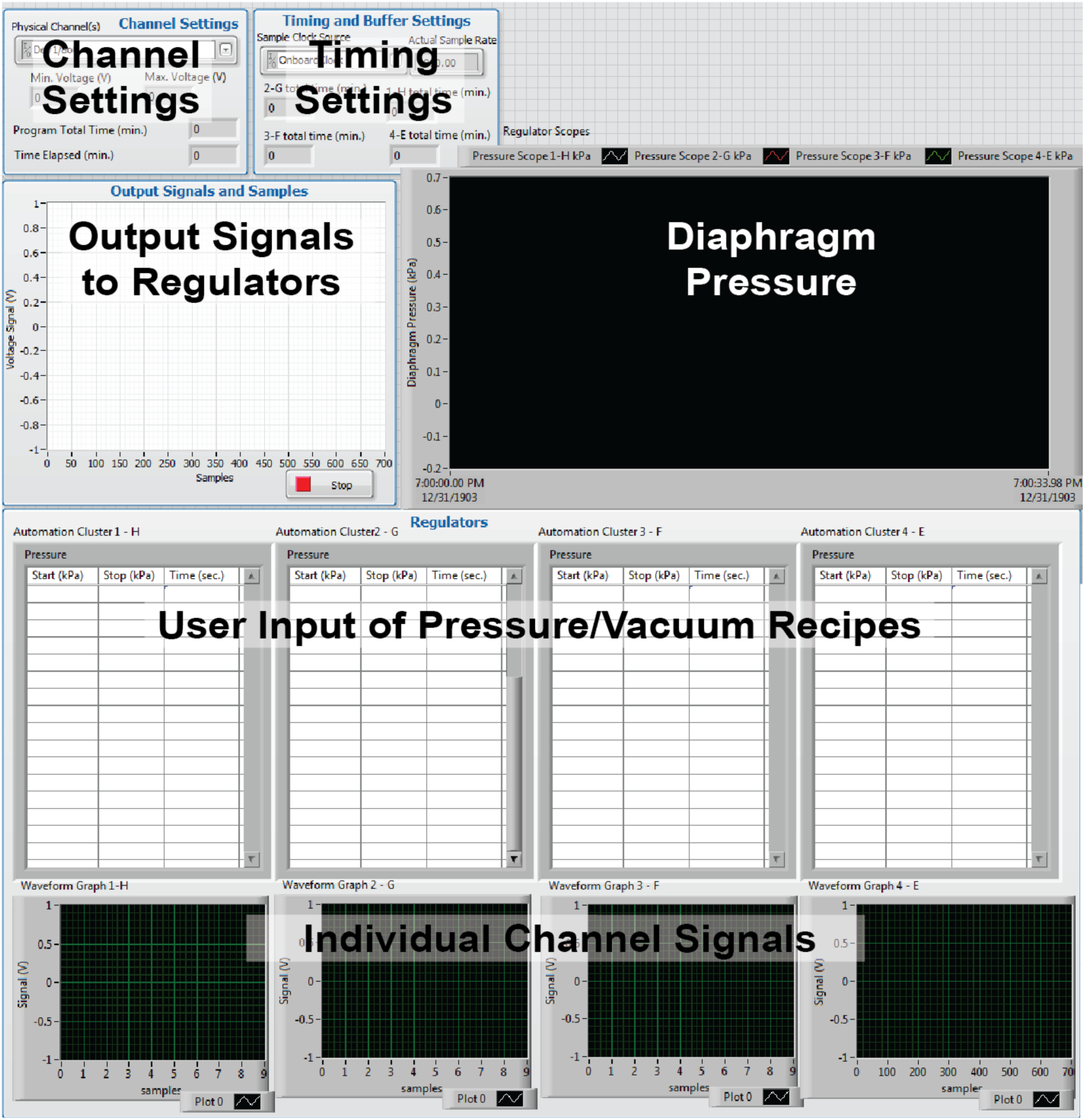
Labview control software. Screenshot of LabVIEW VI used to control the pressure regulators for diaphragm deflections.

### 2. REM-Chip designs

Two REM-Chip geometries were used throughout the study (Fig. S4). The REM-Chip shown in Fig. 2 was used for the majority of experiments. An alternative REM-Chip geometry is shown in Fig. 2. A CAD drawing of the two devices (REMChip.dxf) is available in the online Supplemental Information. In the alternative REM-Chip design, the media line jumps between two layers whereas the main REM-Chip design in Fig. 2 has the media line only in the main PDMS layer. We found that the single layer was simpler to fabricate and operate.

**Figure S4.**
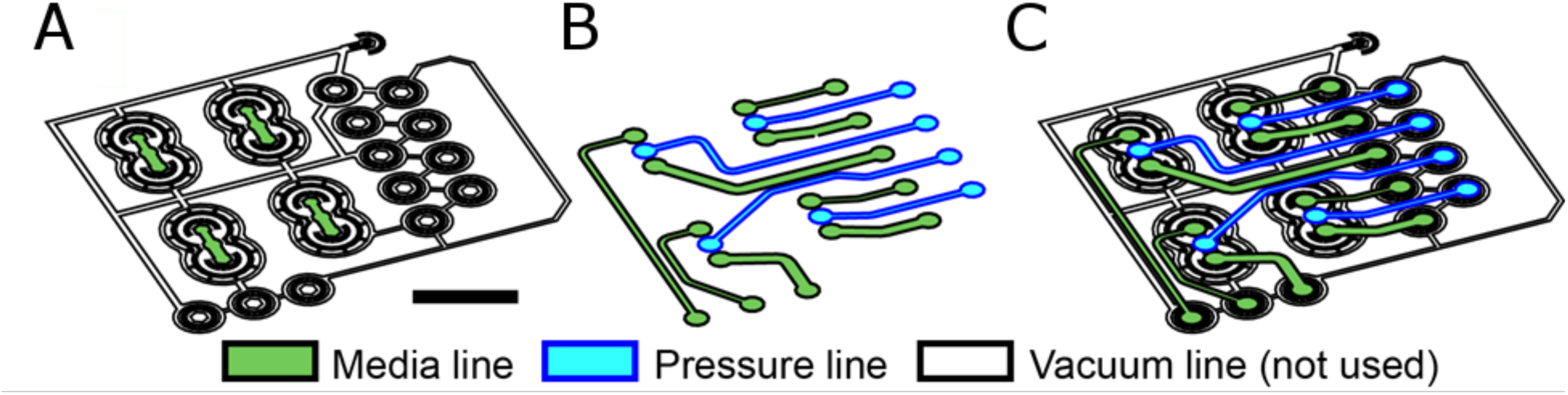
Alternative REM-Chip design used in this study. (a) Bottom device layer with culture chambers in green. The white vacuum lines were not used in this study. (b) Top device layer with media lines (green) and pressure lines. (c) Overlay of top and bottom layers. Scale bar is 5 mm.

### 3. Wing disc loading

Fig. S5 provides a graphical description of the optimal procedure for loading wing discs into the culture chamber. This procedure has proven to be a reliable and efficient means to flow wing discs into the culture chambers.

**Figure S5.**
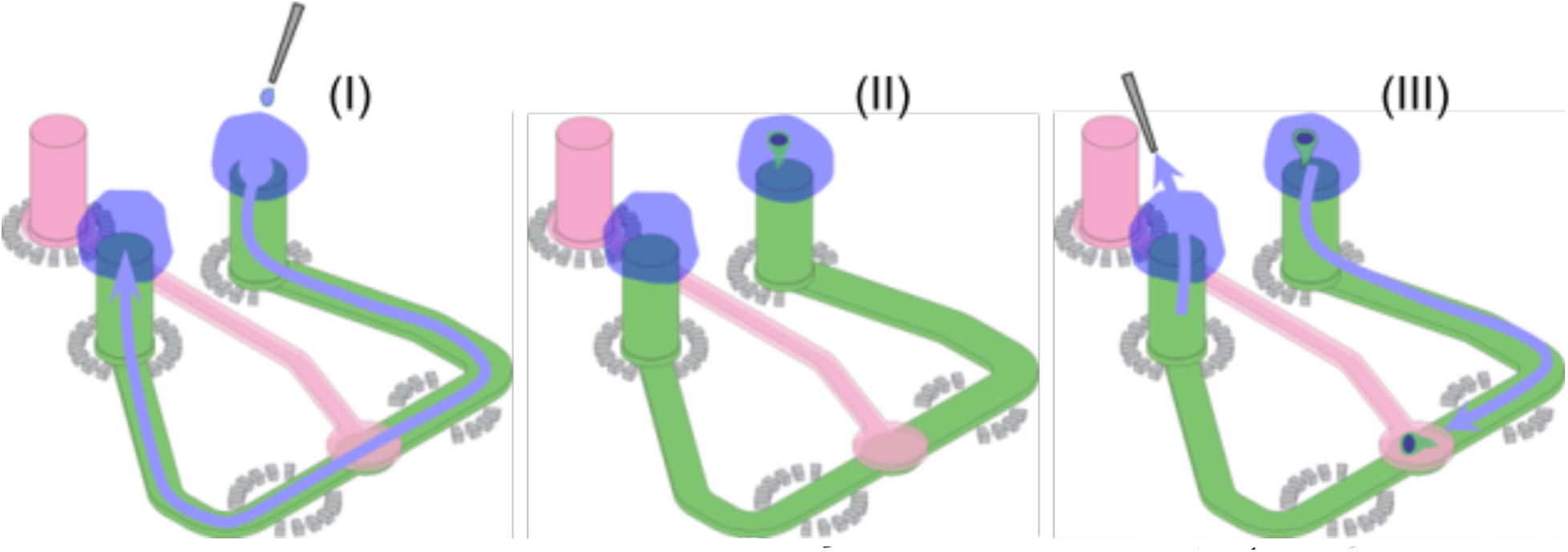
Wing disc loading procedure. The flow channel is first pre-flooded with culture media (I). The wing disc to be loaded is then transferred by pipette to a small bubble of media above the inlet line and positioned such that the dorsal side faces the channel opening (II). The disc is then drawn into the chamber by withdrawing media from the outlet by a pipette (III). At this point, orientation of the disc can be checked and if the pouch is not positioned such that the peripodial membrane is facing the glass coverslip, the disc may be slightly retracted from the chamber by a drawing fluid with a pipette at the inlet and reloaded following the previous procedure until the correct orientation is obtained.

### 4. Diaphragm deflection

#### 4.1 Confocal Imaging and image processing

The PDMS diaphragm was stained with a 0.5 mg/mL Nile red solution in methanol, which was pumped through the chamber at 20 µl/h for 2 h using a syringe pump. During this process, the device and stain-containing syringe were maintained in the dark. The device was then repeatedly flushed at least 3x with DI water until the dye was not observed in the channel’s fluid outlet.

Fluorescent confocal images of the Nile red-stained diaphragm were collected using a 560 nm excitation/640 nm emission filter set, a 10x/0.45 NA objective, and a z-step of 1 µm. Before acquiring each confocal stack, the pressure was allowed to equilibrate for at least 15 s at each set point. Confocal imaging was performed in two fields of view with a 10% overlap, and the resulting images were digitally stitched (1) to create three-dimensional reconstructions of the diaphragm. Cross section A-A’ in Fig. 2a through the center of the diaphragm was then extracted to quantify diaphragm deflection using a custom MATLAB script, which is available for download. This script calculates the centerline of the diaphragms under the deflection conditions studied and outputs spatially scaled deflection profiles based on the imaging parameters.

#### 4.2 Small deflection hysteresis leads to good repeatability

The pressure above the diaphragm was cycled between 5 kPa and 25 kPa (Fig. S6). A small hysteresis is observed between loading and unloading, which is appropriate for this material (2). A small (< 2 µm) increase in the deflection distance was observed with each cycle, indicating the membrane becomes slightly more compliant with use.

**Figure S6.**
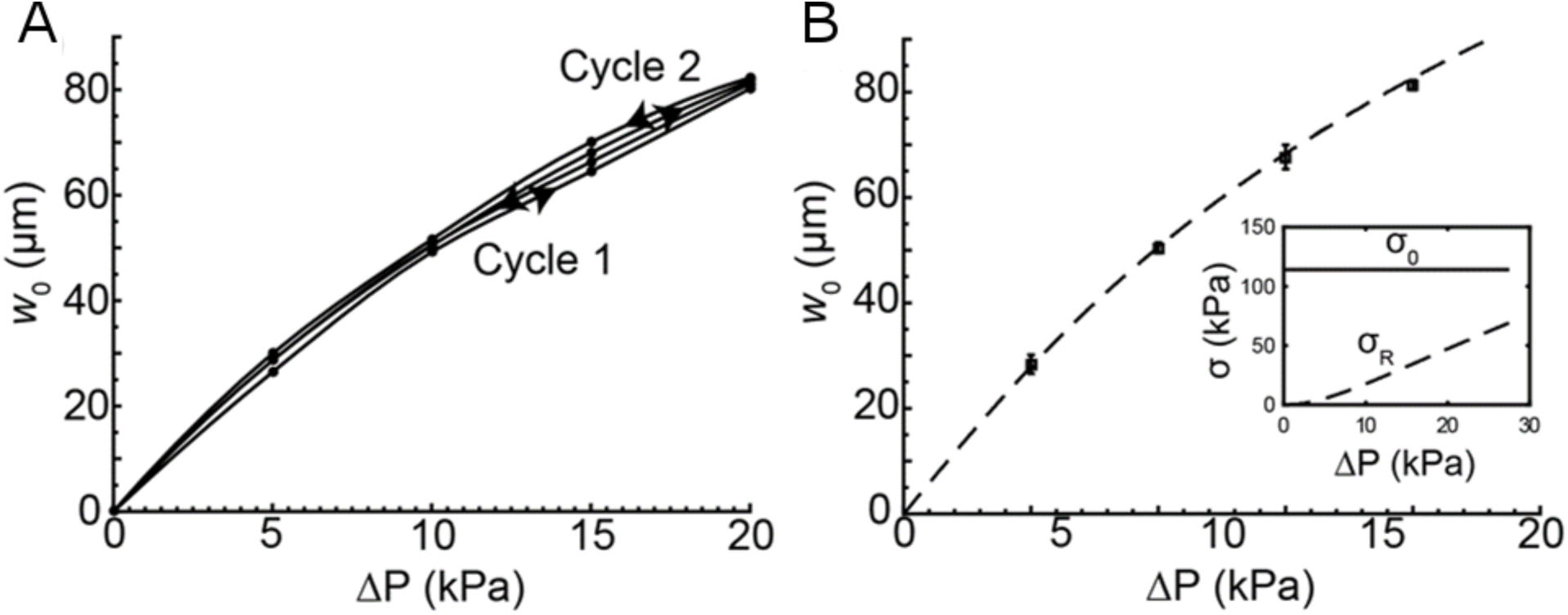
Membrane deflection exhibits minimal hysteresis and agrees with theoretical models. (**a**) Maximum deflection distances during two pressure cycles between 0 kPa and 20 kPa. Arrows indicate cycle direction **(b)** Experimental results and different theoretical solutions for maximum diaphragm deflections at different pressures (inset: Relative contributions of residual stress (*σ*_0_) and stress due to deflection (*σ*_R_) as a function of Δ*P*).

#### 4.3 Theoretical analysis of diaphragm deflection

The measured maximum deflection (*w*_0_) *vs*. the pressure differential across the diaphragm (Δ*P*) is given in Fig. S6b. A general analytical expression for the deflection of a thin diaphragm has been developed based on a semi-empirical fit to finite element method analysis caclulations (3) such that

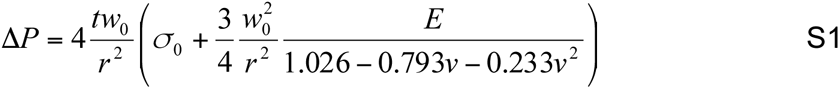

where *t* is the diaphragm thickness (~76 µm), *r* is diaphragm’s radius (400 µm), *E* is the Young’s modulus of PDMS (1200 kPa for ~100 µm-thick PDMS film (4)), *v* is the Poisson ratio of PDMS (0.5), and *σ*_0_ is the diaphragm’s residual stress. The results were fit to Equation S1 (line in Fig. S6b) using *σ*_0_ = 116 kPa, which is a reasonable value for a thin PDMS film (5). The right-hand component of Equation S1 is the stress due to diaphragm deflection, *σ*_R_. The contributions of the two stress components are shown as a function of Δ*P* in the inset to Fig. S6b.

### 5. Fly lines and reagents

General fly lines used in the present study:

1. <u>P{UAS-Dcr-2.D}1</u>, w^1118^; <u>P{GawB}nubbin-AC-62</u> (BL#25754) – nubbin-GAL4 line
2. w^1118^ P{20XUAS-IVS-GCaMP6f}attP40 (BL#42747) – UAS-GCaMP6f line
3. w^1118^; GCaMP6f, nubbin-GAL4/CyO - tester line created by recombining BL#25754 with BL#42747. This enabled single crosses to RNAi lines as described in the text.
4. DECadherin::GFP(6). E-Cadherin::GFP provides a marker of the apical surface of wing discs.
5. Lac::YFP (KSC# CPTI – 002601 and described in (7)). Lac::YFP is a transgenic line providing a marker for the septate junctions.

RNAi Lines:

6. BL#25937: *UAS-IP_3_R^RNAi^* (documented wound healing phenotype in (8))
7. BL#29306: UAS-*Inx2^RNAi^*
8. BL#27263: SERCA^RNAi^

#### Abbreviations

BL: Bloomington Stock Center
KSC: Kyoto Stock Center

Reagents used:

Schneider’s Drosophila Media – Gibco (21720-024)

Human Insulin – Sigma Aldrich (I9278)

Penicillin-Streptomycin – Gibco (15140-122)

Phosphate buffered saline – Sigma Aldrich (D5652)

ZB media was prepared in house as describe in (9)

Fig. S7 shows representative phenotypes of RNAi crossed to the tester line.

**Figure S7.**
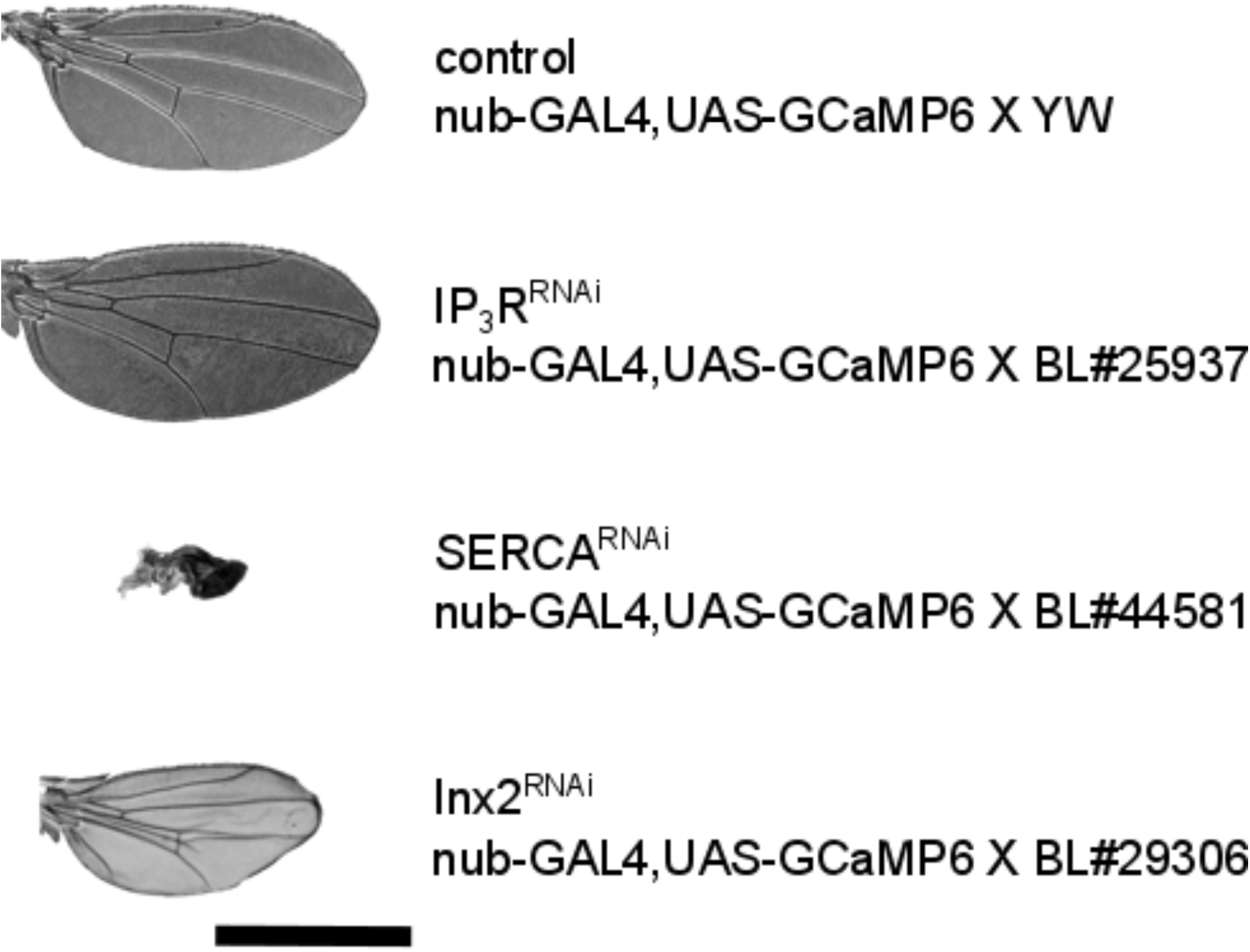
Wing phenotypes of the RNAi crosses. Shown above are representative wing phenotypes for each of the RNAi lines tested. Progeny from the tester line nub-Gal4, UAS-GCaMP6f crossed to yw show normal wings. Although *IP_3_R* does not show a dramatic developmental phenotype, this BL line has been previously validated with a wound healing phenotype in(8) and results in larger wings compared to the control *SERCA*^RNAi^ leads to crumpled wings and *inx2* shows a darkening of the wing, vein defects and smaller size compared to the control. Scale bar is 2 mm.

### 6. Quantification methods and additional supplemental studies

ICW velocities were manually extracted from images by measuring the distance of travel over a known time (Fig. S8a). Delays between release of compression and IWC initiation and burst times were determined by visually monitoring the onset and completion of ICWs. The burst area, or percent of the pouch the ICW covered, was extracted using a custom MATLAB script (“burst_area.m”) (Fig. S8b). In this script, waves were quantified in pre-specified image frames (corresponding to initial and final times of *t* = 1 and *t* = final, respectively) by calculating the 1-norm of the change in pixel intensities between adjacent scripts according to

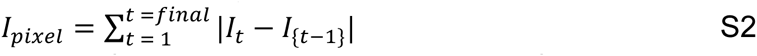

where *I_pixel_* is the ICW of intensity of a pixel, *I_t_* is the intensity in the frame corresponding to time *t*, and *I_t_*_-1_ is the intensity of the pixel in the preceding frame. Images were reconstructed using the values of *I_pixel_* to map ICWs after three sequential compressions, as shown in Fig. S8. These images were binarized, and the total number of pixels within the waves were counted. Burst area was calculated as

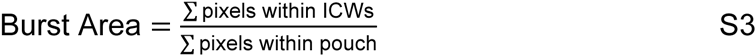

**Figure S8.**
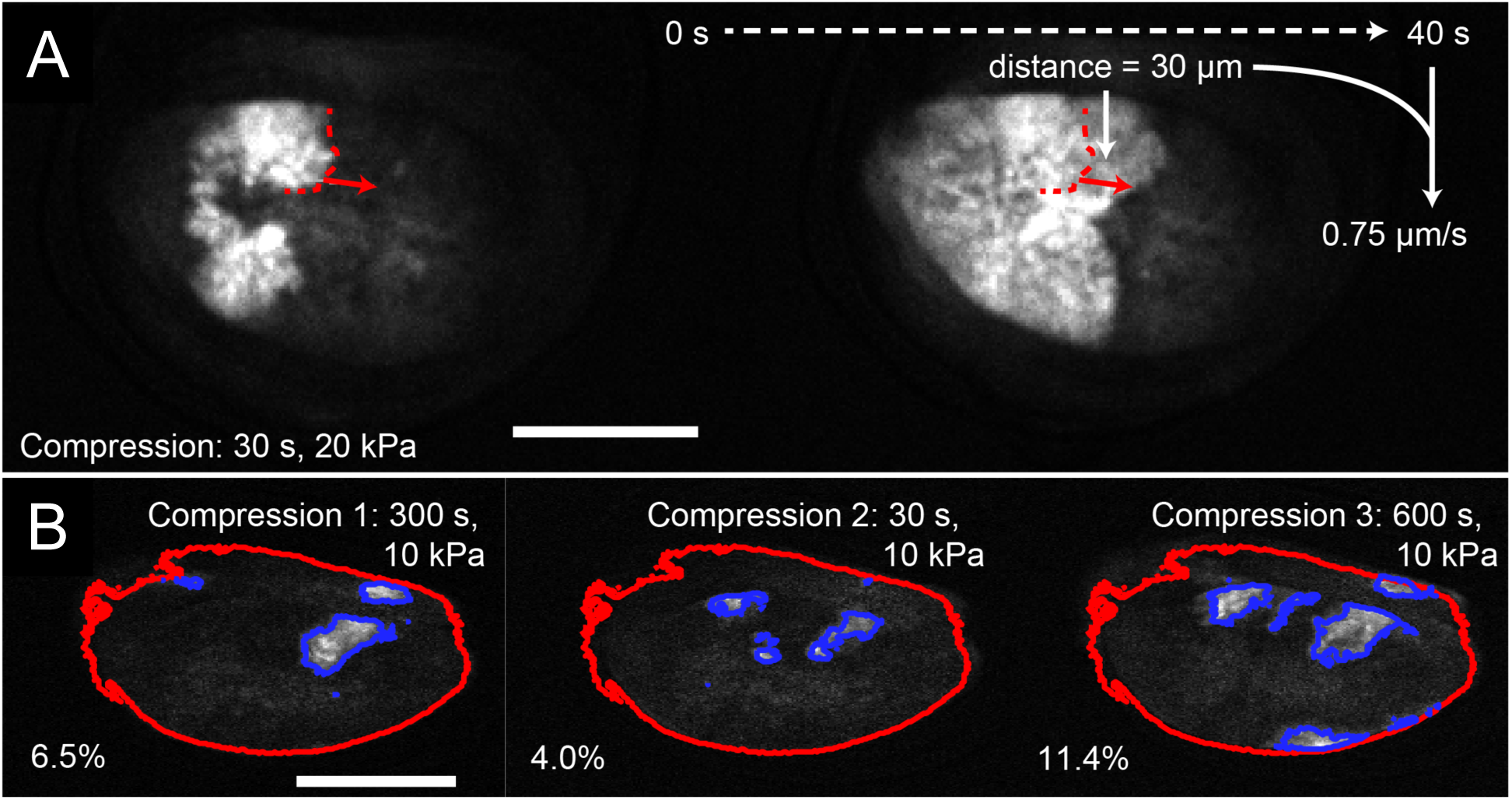
Calculation of ICW velocity and burst area. (**a**) Depiction of velocity calculation procedure for a representative case. The distance the front (dashed red line) of a selected ICW travels (red arrow, 30 µm) over a known time (40 s) was used calculate wave velocity (0.75 µm/s). Scale bar is 100 µm. (**b**) Output of MATLAB script (burst_area.m) used to calculate the percentage of the pouch area covered by ICWs. Red line is an outline of the pouch, and blue outlined regions are ICWs. All scale bars are 100 µm.

Peak mean pouch intensity following an ICW was obtained using ImageJ to extract the mean pouch intensity. The pouch was outlined as a region of interest (ROI) in each frame by duplicating the confocal video stack and using Phansalkar auto local thresholding (Image>Adjust>Auto Local Threshold) to create a binary mask. The “Analyze Particles…” function was used to add the pouch mask as an ROI in each frame and the disc area and mean intensity extracted. The mean intensity value was normalized to the basal Ca^2+^ in the pouch, which was estimated by averaging 5 frames for each video that did not contain an ICW. The max value in the 30 frames after each compression event was then taken as the normalized peak mean pouch intensity. This method was able to provide masks and mean intensity measurements for 70% of experimental videos (*n* = 13/19 wing disc videos with ICWs)

To classify the baseline activity of each wing disc, confocal image stacks were first randomized using the file name randomizer described above and opened in ImageJ. Activity was qualitatively scored as “low” or “high” depending on the baseline activity of the disc prior to compression during the first ten minutes of each experiment. Low activity discs demonstrated no activity, minimal discrete cell flashing, or very small infrequent transients. High activity discs demonstrated frequent abundant small transients and/or one or more large ICWs before the first mechanical perturbation event.

**Figure S9.**
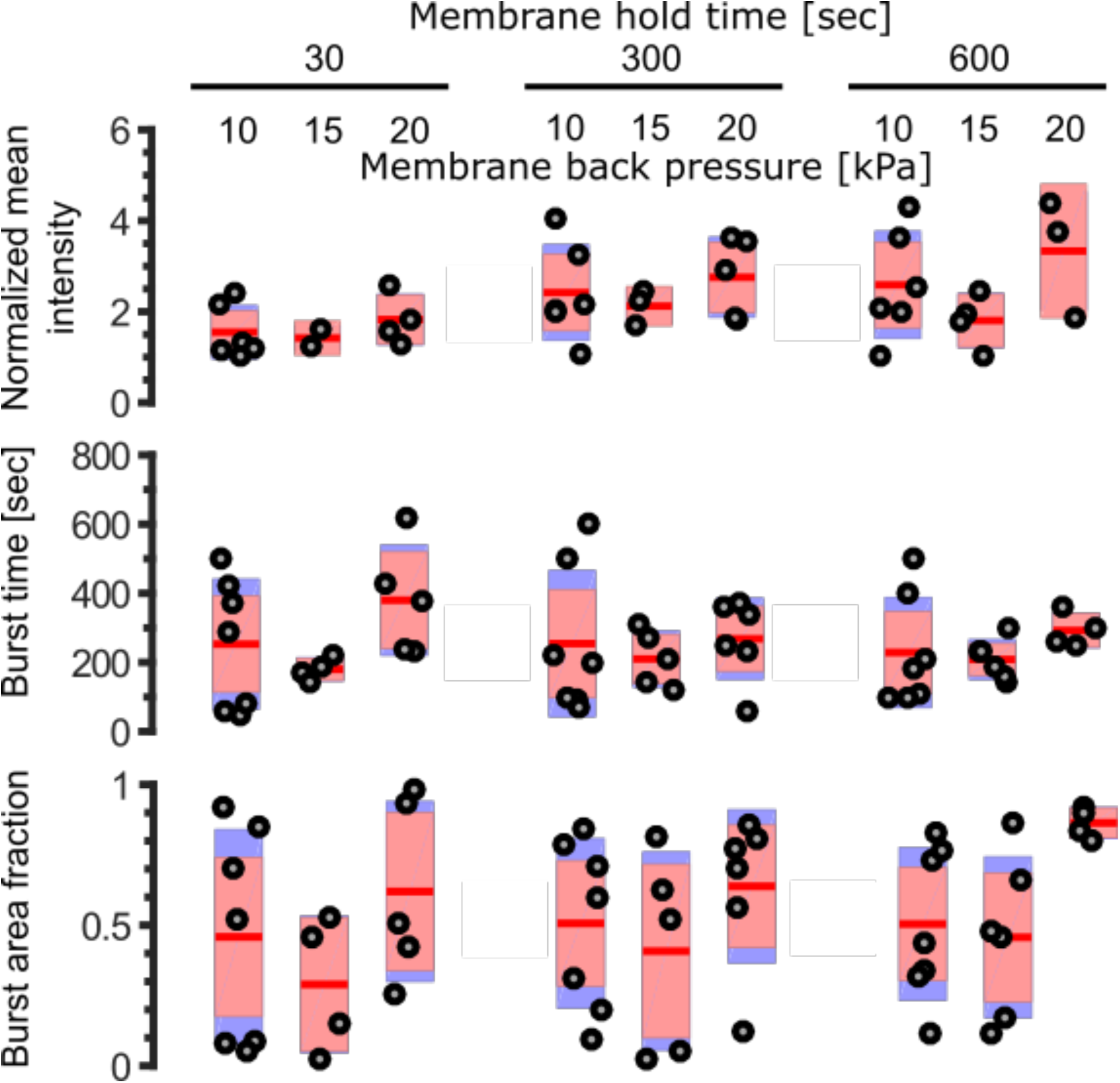
ICW properties do not show significant dependence on applied backpressure or hold time. (Top) Plot of normalized peak mean pouch intensity with respect to applied backpressure and hold time. (Middle) Plot of burst time with respect with respect to applied backpressure and hold time. (Bottom) Plot of burst area as a fraction of pouch area with respect to applied backpressure and hold time. A general trend can be observed in the effects of applied back pressure and hold time on burst time and normalized mean intensity.

### 7. Description of SI movies

#### List of SI Movies

SI Movie 1: Fig. 1b Representative movie of *in vivo* waves

SI Movie 2: Fig. 1c Representative movie of diaphragm deflection

SI Movie 3: Fig. 4c Representative movie of a disc in ZB media

SI Movie 4: Fig. 4d Representative movie of a disc in ZB media + 15% FEX SI Movie 5: Fig. 5c Control

SI Movie 6: Fig. 5c IP_3_R^RNAi^

SI Movie 7: Fig. 5c SERCA^RNAi^ SI Movie 8: Fig. 5c inx2^RNAi^

SI Movie 9: Representative movie of ICW immediately before application of compression

### 8. Description of additional SI files

REM-Chip designs: REM-Chip.dxf

Code for diaphragm deflection: diaphragm_deflection.zip

VI code for pressure regulator: pressure_automation.vi

File name randomizer: RandomNames.zip

Burst Area Analysis: burst_area.zip

## REFERENCES

1. Berridge MJ, Lipp P, Bootman MD (2000) The versatility and universality of calcium signalling. Nat Rev Mol Cell Biol 1(1):11–21.

2. Lebeche D, Davidoff AJ, Hajjar RJ (2008) Interplay between impaired calcium regulation and insulin signaling abnormalities in diabetic cardiomyopathy. Nat Rev Cardiol 5(11):715–724.

3. Prevarskaya N, Skryma R, Shuba Y (2011) Calcium in tumour metastasis: new roles for known actors. Nat Rev Cancer 11(8):609–618.

4. Berridge MJ (2013) Dysregulation of neural calcium signaling in Alzheimer disease, bipolar disorder and schizophrenia. Prion 7(1):2–13.

5. Deng H, Gerencser AA, Jasper H (2015) Signal integration by Ca2+ regulates intestinal stem-cell activity. Nature 528(7581):212–217.

6. Antunes M, Pereira T, Cordeiro JV, Almeida L, Jacinto A (2013) Coordinated waves of actomyosin flow and apical cell constriction immediately after wounding. J Cell Biol 202(2):365–379.

7. Restrepo S, Basler K (2016) Drosophila wing imaginal discs respond to mechanical injury via slow InsP3R-mediated intercellular calcium waves. Nat Commun 7:12450.

8. Renò F, et al. (2003) In vitro mechanical compression induces apoptosis and regulates cytokines release in hypertrophic scars. Wound Repair Regen 11(5):331–336.

9. Hariharan IK (2015) Organ Size Control: Lessons from Drosophila. Dev Cell 34(3):255–265.

10. Shraiman BI (2005) Mechanical feedback as a possible regulator of tissue growth. Proc Natl Acad Sci U S A 102(9):3318–3323.

11. Chiou KK, Hufnagel L, Shraiman BI (2012) Mechanical Stress Inference for Two Dimensional Cell Arrays. PLOS Comput Biol 8(5):e1002512.

12. Schluck T, Nienhaus U, Aegerter-Wilmsen T, Aegerter CM (2013) Mechanical Control of Organ Size in the Development of the Drosophila Wing Disc. PLoS ONE 8(10):e76171.

13. Yang S, Saif T (2005) Micromachined force sensors for the study of cell mechanics. Rev Sci Instrum 76(4):44301.

14. Zhang X, Dhumpa R, Roper MG (2013) Maintaining stimulant waveforms in largevolume microfluidic cell chambers. Microfluid Nanofluidics 15(1):65–71.

15. Lucchetta EM, Lee JH, Fu LA, Patel NH, Ismagilov RF (2005) Dynamics of Drosophila embryonic patterning network perturbed in space and time using microfluidics. Nature 434(7037):1134–1138.

16. Bleuel J, Zaucke F, Brüggemann G-P, Niehoff A (2015) Effects of Cyclic Tensile Strain on Chondrocyte Metabolism: A Systematic Review. PLOS ONE 10(3):e0119816.

17. Fatehullah A, Tan SH, Barker N (2016) Organoids as an in vitro model of human development and disease. Nat Cell Biol 18(3):246–254.

18. Gilbert LI (2008) Drosophila is an inclusive model for human diseases, growth and development. Mol Cell Endocrinol 293(1–2):25–31.

19. Schomburg PD rer nat WK (2011) Membranes. Introduction to Microsystem Design, RWTHedition. (Springer Berlin Heidelberg), pp 29–52.

20. Chiou C-H, Yeh T-Y, Lin J-L (2015) Deformation Analysis of a Pneumatically-Activated Polydimethylsiloxane (PDMS) Membrane and Potential Micro-Pump Applications. Micromachines 6(2):216–229.

21. Zartman J, Restrepo S, Basler K (2013) A high-throughput template for optimizing Drosophila organ culture with response-surface methods. Dev Camb Engl 140(3):667–674.

22. Restrepo S, Zartman J, Basler K (2016) Cultivation and Live Imaging of Drosophila Imaginal Discs. Drosophila, Methods in Molecular Biology., ed Dahmann C (Springer New York), pp 203–213.

23. Burnette M, Brito-Robinson T, Li J, Zartman J (2014) An inverse small molecule screen to design a chemically defined medium supporting long-term growth of Drosophila cell lines. Mol Biosyst. doi:10.1039/C4MB00155A.

24. Narciso C, et al. (2015) Patterning of wound-induced intercellular Ca 2+ flashes in a developing epithelium. Phys Biol 12(5):56005.

25. Razzell W, Evans IR, Martin P, Wood W (2013) Calcium flashes orchestrate the wound inflammatory response through DUOX activation and hydrogen peroxide release. Curr Biol CB 23(5):424–429.

26. Kaneuchi T, et al. (2015) Calcium waves occur as Drosophila oocytes activate. Proc Natl Acad Sci 112(3):791–796.

27. Mammoto T, Mammoto A, Ingber DE (2013) Mechanobiology and Developmental Control. Annu Rev Cell Dev Biol 29(1):27–61.

28. Buchmann A, Alber M, Zartman JJ (2014) Sizing it up: The mechanical feedback hypothesis of organ growth regulation. Semin Cell Dev Biol 35C:73–81.

29. Farge E (2003) Mechanical induction of Twist in the Drosophila foregut/stomodeal primordium. Curr Biol CB 13(16):1365–1377.

30. Ghannad-Rezaie M, Wang X, Mishra B, Collins C, Chronis N (2012) Microfluidic Chips for In Vivo Imaging of Cellular Responses to Neural Injury in Drosophila Larvae. PLoS ONE 7(1):e29869.

31. Huh D, et al. (2010) Reconstituting organ-level lung functions on a chip. Science 328(5986):1662–1668.

32. Dhumpa R, Truong TM, Wang X, Bertram R, Roper MG (2014) Negative feedback synchronizes islets of Langerhans. Biophys J 106(10):2275–2282.

33. Xia Y, Whitesides GM (1998) Soft Lithography. Angew Chem Int Ed 37(5):550–575.

34. Lake M, et al. (2015) Microfluidic device design, fabrication, and testing protocols. Protoc Exch. doi:10.1038/protex.2015.069.

35. Unger MA, Chou H-P, Thorsen T, Scherer A, Quake SR (2000) Monolithic Microfabricated Valves and Pumps by Multilayer Soft Lithography. Science 288(5463):113–116.

36. Schindelin J, et al. (2012) Fiji: an open-source platform for biological-image analysis. Nat Methods 9(7):676–682.

37. Huang J, Zhou W, Dong W, Watson AM, Hong Y (2009) Directed, Efficient, and Versatile Modifications of the Drosophila Genome by Genomic Engineering. Proc Natl Acad Sci 106(20):8284–8289.

38. Duffy JB (2002) GAL4 system in drosophila: A fly geneticist’s swiss army knife. genesis 34(1–2):1–15.

39. Tian L, et al. (2009) Imaging neural activity in worms, flies and mice with improved GCaMP calcium indicators. Nat Methods 6(12):875–881.

40. Ashburner M (1989) Drosophila: A laboratory handbook (Cold Spring Harbor Laboratory).

## Supplemental references

1. Preibisch S, Saalfeld S, Tomancak P (2009) Globally optimal stitching of tiled 3D microscopic image acquisitions. Bioinformatics 25(11):1463–1465.

2. Selby JC, Shannon MA (2007) Apparatus for measuring the finite load-deformation behavior of a sheet of epithelial cells cultured on a mesoscopic freestanding elastomer membrane. Rev Sci Instrum 78(9):94301.

3. Schomburg WK (2015) Introduction to Microsystem Design (Springer).

4. Liu M, Sun J, Sun Y, Bock C, Chen Q (2009) Thickness-dependent mechanical properties of polydimethylsiloxane membranes. J Micromechanics Microengineering 19(3):35028.

5. Thangawng AL, Ruoff RS, Swartz MA, Glucksberg MR (2007) An ultra-thin PDMS membrane as a bio/micro–nano interface: fabrication and characterization. Biomed Microdevices 9(4):587–595.

6. Huang J, Zhou W, Dong W, Watson AM, Hong Y (2009) Directed, efficient, and versatile modifications of the Drosophila genome by genomic engineering. Proc Natl Acad Sci:pnas.0900641106.

7. Zartman J, Restrepo S, Basler K (2013) A high-throughput template for optimizing Drosophila organ culture with response-surface methods. Dev Camb Engl 140(3):667–674.

8. Restrepo S, Basler K (2016) Drosophila wing imaginal discs respond to mechanical injury via slow InsP3R-mediated intercellular calcium waves. Nat Commun 7:12450.

9. Burnette M, Brito-Robinson T, Li J, Zartman J (2014) An inverse small molecule screen to design a chemically defined medium supporting long-term growth of Drosophila cell lines. Mol Biosyst. doi:10.1039/C4MB00155A.

